# Exosomes derived from highly scalable and regenerative human progenitor cells promote functional improvement in a rat model of ischemic stroke

**DOI:** 10.1101/2025.01.07.631793

**Authors:** Jieun Lee, Susanna R. Var, Derek Chen, Dilmareth E. Natera-Rodriguez, Mohammad Hassanipour, Michael D. West, Walter C. Low, Andrew W. Grande, Dana Larocca

## Abstract

Globally, there are 15 million stroke patients each year who have significant neurological deficits. Today, there are no treatments that directly address these deficits. With demographics shifting to an older population, the problem is worsening. Therefore, it is crucial to develop feasible therapeutic treatments for stroke. In this study, we tested exosomes derived from embryonic endothelial progenitor cells (eEPC) to assess their therapeutic efficacy in a rat model of ischemic stroke. Importantly, we have developed purification methods aimed at producing robust and scalable exosomes suitable for manufacturing clinical grade therapeutic exosomes. We characterized exosome cargos including RNA-seq, miRNAs targets, and proteomic mass spectrometry analysis, and we found that eEPC-exosomes were enhanced with angiogenic miRNAs (i.e., miR-126), anti-inflammatory miRNA (i.e., miR-146), and anti-apoptotic miRNAs (i.e., miR-21). The angiogenic activity of diverse eEPC-exosomes sourced from a panel of eEPC production lines was assessed *in vitro* by live-cell vascular tube formation and scratch wound assays, showing that several eEPC-exosomes promoted the proliferation, tube formation, and migration in endothelial cells. We further applied the exosomes systemically in a rat middle cerebral artery occlusion (MCAO) model of stroke and tested for neurological recovery (mNSS) after injury in ischemic animals. The mNSS scores revealed that recovery of sensorimotor functioning in ischemic MCAO rats increased significantly after intravenous administration of eEPC-exosomes and outpaced recovery obtained through treatment with umbilical cord stem cells. Finally, we investigated the potential mechanism of eEPC-exosomes in mitigating ischemic stroke injury and inflammation by the expression of neuronal, endothelial, and inflammatory markers. Taken together, these data support the finding that eEPCs provide a valuable source of exosomes for developing scalable therapeutic products and therapies for stroke and other ischemic diseases.

## Introduction

Stroke continues to be the primary cause of morbidity in the United States (US), with about 795,000 cases annually costing over $53 billion per year^1,2^. To date, there remains just one drug approved for stroke intervention, tissue plasminogen activator (tPA), but the use of tPA is limited due to its narrow therapeutic time window of 6 to 8 hours from the onset of ischemic stroke. Thus, most patients are not able to receive tPA administration in a timely manner, which results in fewer than 5% of stroke patients ever benefiting from the needed treatment ^3^. Stroke patients with a higher cerebral blood vessel density in the cortical infarct rim show better progress and survival than those who have lower levels of vascular density ^4–6^. Preclinical studies have shown that angiogenesis promotes ischemic brain repair, and that blockage of cerebral angiogenesis inhibits stroke recovery ^7–9^. However, stroke-increased angiogenesis is limited, thus minimizing functional recovery ^7–9^.

Recognizing the importance of finding therapies that can be used beyond the 8-hour time window after stroke, therapies using stem cell have become promising candidates for the treatment of ischemic stroke. These methods use different cell sources, including neural stem cells (NSCs) ^10–12^, pluripotent stem cells (PSCs) ^13,14^, mesenchymal stem cells (MSCs) ^18–20^, umbilical cord blood stem cells (UCBSCs) ^8,15–17^, and adipose-derived stem cells (ADSCs) ^21,22^. Clinical trials utilizing cell transplantation (i.e., BMSCs, MSCs, and UCSCs) have shown promising results ^23–25^, yet the effectiveness of stem cell therapy in treating stroke has not yet been confirmed in clinical trials ^26,27^. The primary purpose of stem cell administration after stroke has been to regenerate neurons that are able to integrate into host tissue in order to replace lost or damaged neurons. However, a great amount of evidence has shown that systemically delivered mesenchymal stem cells (MSCs) become trapped within the lung ^10,28^, resulting in low or even undetectable cell numbers in the ischemic brain. This indicates that the therapeutic effects of stem cells may be in large part due to paracrine factors through interaction with brain parenchymal cells, which can impact neuroprotection, immune modulation, and neovascular remodeling. In this regard, reports have indicated that treatment with cultured MSC supernatant not only improved the function of endothelial cells, but also brought in new macrophages to bolster wound healing in ischemia ^18,29^. This indicated that MSCs secreted multiple growth factors and cytokines which include angiogenetic factors ^31^, neuroprotective factors ^30^, and anti-inflammatory cytokines ^32, 33,34^. More recently, extracellular nanoparticle-sized vesicles known as exosomes have been isolated from conditioned medium of cultured stem cells, thus supporting the finding that exosomes, secreted from stem cells, enhance cellular communication between brain parenchymal cells and stem cells, thus promoting therapeutic effects ^30,35–38^. Importantly, exosomes are known to have an ability to internalize into recipient cells by passing through the blood–brain barrier (BBB) ^39^ and into brain parenchyma ^15,40,41^. We therefore propose here to use stem cell derived exosomes as a means of promoting neurovascular remodeling and neurological recovery after stroke.

Exosomes originate from the endosomal membrane compartment ^42^ and are contained inside intraluminal vesicles located within multivesicular bodies of the late endosome. Multivesicular bodies come from the early endosome compartment and possess smaller vesicular bodies inside, including exosomes ^43^. Exosomes exit a cell when the plasma membrane unites with multivesicular bodies (MVBs). Exosomes are identifiable by certain markers of biochemical composition (i.e., CD63, CD81, CD9, Tsg101 and Alix) ^44,45^ along with physical characteristics such as size. For instance, “small EVs” are less than 150nm in diameter ^46,47^. Recent studies have shown that exosomes function as biological transporters ^42^ that transfer proteins or genetic materials to promote cell repair ^31,48^, boost angiogenesis ^31,49,50^, or even have regulatory effect on inflammation ^49^, which demonstrates the potential therapeutic application of exosomes for treatment of cardiovascular disease ^9,49,51^.

Various adult stem cells such as mesenchymal stem cells (MSCs) ^26,54^, endothelial progenitor cells (EPCs) ^52,53^, and cardiosphere-derived cells (CDCs) ^55^ are presently being tested for their potential as therapeutic agents ^52^ as a source of exosomes for ischemic diseases like stroke. However, the effective use of adult stem cell therapies for treating a large patient population is made challenging ^56^ due to issues of identification, stability, scalability, and purity. We previously reported a process for developing pure embryonic site-specific somatic cells by propagating hPSC-derived clonal human embryonic progenitor cell lines ^57^ that can mitigate current limitations of purity and scale. We identified these cultures to be human “embryonic progenitor” (hEP) cells ^58^ based on their ability to autonomously renew, their strong expression of embryonic developmental stage gene markers, and the fact that they lack fetal/adult gene markers that are typically expressed in cells that have crossed the embryonic-fetal transition ^59^. We specifically derived embryonic endothelial progenitor cell (eEPC) lines possessing endothelial properties to be a source of human endothelial cells and cell products that are scalable for therapeutics in ischemic disease. We found that exosomes secreted from eEPCs exhibit enriched angiogenic factors such as miR-126 ^60–62^. This demonstrated the potential for highly scalable eEPCs to generate exosomes with greater angiogenic potency compared to bone marrow-derived MSCs (BM-MSC) ^58^.

In the present study, we 1) developed an optimized exosome purification method, enabling subsequent downstream therapeutic biomarkers that are stable and effective, 2) identified production cell lines that produce eEPC-exosomes with high angiogenic activity *in vitro* and characterized their cargo contents, which included angiogenic microRNAs and proteins, and 3) demonstrated that *in vitro* angiogenic eEPC-exosomes were effective *in-vivo* in the treatment of stroke using a rat middle cerebral artery occlusion (MCAO) model.

## Materials and Methods

### Human embryonic endothelial progenitor cell (eEPC) culture and exosome collection

As previously reported, human eEPCs were cultured in endothelial growth medium (EGM-MV2, Cat# C-39226, Lot# 25 x 461M089, PromoCell, GmbH, Germany) on gelatin-coated plates. The medium was changed every 2-3 days and cells were passaged at 80% confluency ^58^. The cells when maintained in the undifferentiated state were cultured at 37°C in a humidified atmosphere of 10% CO_2_ and 5% O_2_. The medium was changed every 2-3 days and cells were passaged at 80-90% confluence with TrypLE Express medium. For exosome collection, eEPCs between passages 10 and 13 were used. After cells reached ∼80% confluence, they were washed two times with PBS. Medium was changed with medium containing endothelial basal medium (EBM) supplement with VEGF, IGF and FGF, cultures were incubated for 72 hours at 5% oxygen, and conditioned media were collected for exosome purification.

### Exosome purification

Conditioned media were centrifuged at 300g for 5 min followed by 1000g for 10 min at room temperature and filtered through 0.2 µm to remove cells and cellular debris. Conditioned medium was then subjected to ultrafiltration in Tangential Flow Filtration (TFF) system using a 100 kDa cutoff TFF cartridge (PALL Laboratory, New York). A feed flow rate of 40mL/min with transmembrane pressure <2 psi was applied. The conditioned medium was concentrated 10-fold and centrifuged at 10,000g for 10 min. Continuously, size exclusion chromatography (SEC) using qEV100 columns (Izon Science, Cambrige, MA) was performed for further purification of exosomes. Briefly, after rinsing the qEV columns with PBS, 100ml of TFF-concentrated exosomes were eluted with 6 fractions (F1-F6, total 150ml). A total of F1-F6 fractions were pooled and further concentrated. Amicon Ultrafilter-70 Centrifugal Filters (100KDa MWCO, Millipore, MA) were used to concentrate exosomes. Purified exosomes were aliquoted with 100 µL and stored at -80°C.

### Nanoparticle tracking analysis

According to the manufacturer’s software manual (NanoSight), exosome subpopulations were analyzed using NanoSight LM10 (Malvern Analytical), to determine particle size and concentration. All samples were diluted 1:100 in PBS prior to analysis. Five measurements of 30 s each were taken with consistent acquisition settings (gain = 2.3, camera level = 14). Data analysis was performed using NTA 3.1 software (NanoSight, Malvern Analytical) with consistent analysis settings (gain = 3, detection threshold = 4). Median particle size, concentration, and size distribution were obtained for all exosome subpopulations.

### Tunable Resistive Pulse Sensing

The size distribution and particle concentration of exosomes were measured using the Tunable Resistive Pulse Sensing (TRPS) qNano platform (iZON® Science, UK). The instrument was set up and calibrated as per manufacturer recommendations. A polyurethane nanopore (NP150, Izon Science) was used and axially stretched to 47 mm, as measured on the qNano unit. Data processing and analysis were carried out on Izon Control Suite software v3.3 (Izon Science).

### Exosome protein measurement

The purified exosomes were resuspended in 100 µL of PBS and lysed in RIPA buffer. Protein quantity was measured by a bicinchoninic acid (BCA) assay using the Micro BCA Protein Assay Kit (Thermo) according to the manufacturer’s instructions. Exosome protein content was determined by calibration against a standard curve, which was prepared by plotting the absorbance at 562 nm versus BSA standard concentration.

### Live cell wound healing migration assay

Cell migration was assessed using a scratch wound healing assay format. HUVEC (1E4 cells per well) were plated onto 0.1% gelatin coated 96-well plates, and the following day a scratch was made on confluent monolayers using a 96-pin WoundMaker (Essen BioScience, Ann Arbor, MI). Exosomes (1E7, 2E7, 4E7 and 1.2E8 particles per well) and growth factor (4 ng/ml VEGF as a positive control) were treated with exosome-depleted EGM-MV2. Wound images were automatically acquired by the IncuCyte software system every 2 hours for 24 hours. Wound closure and cell confluence were calculated using the IncuCyte 96-Well Cell Migration Software Application Module. Migration data were analyzed as the Percent of Relative Wound Density (% RWD). RWD is a representation of the spatial cell density in the wound area relative to the spatial cell density outside of the wound area at every time point (time-curve).

### Angiogenesis tube formation assay

CellPlayer Angiogenesis PrimeKit (Essen BioScience) was performed according to the manufacture’s protocol. On day 0, normal human dermal fibroblasts (NHDFs) were plated into a 96-well plate and then incubated at room temperature in a tissue culture hood for 1 hour to allow them to adhere to the plate. The HUVEC-CytoLight Green were then plated onto the NHDF feeder layers and incubated at room temperature for 1 hour prior to placing in the IncuCyte (Essen BioScience) for imaging. The next day, treatment initiated with a media change including exosomes (4E7 particles per well) and growth factor (4 ng/ml VEGF as a positive control) in exosome-depleted EGM-MV2. Cultures were then fed every 3 days at which time complete media changes occurred with fresh growth factor and the addition of exosomes. Following seeding, co-cultures were placed in an IncuCyte live imaging system, and images were automatically acquired in both phase and fluorescence every 6 hours for 10 to 14 days at 10X objective magnification using the tiled field of view mosaic imaging mode. In this mode, six images were acquired per well and merged into a single larger image. Tube formation over the 14 days was quantified using the IncuCyte Angiogenesis Analysis Module. For analyzing angiogenesis, we used the metric of tube network length (mm/mm^2^) by measuring lengths of all of the networks in the image divided by the image area at every time point.

### Small RNA-sequencing Analysis

Total RNA from exosomes was isolated using the Norgen Total RNA Kit (Norgen Biotek, Cat. # 58000) according to the manufacturer’s instructions. Briefly, exosome samples were lysed and denatured using Lysis Buffer A and Lysis Additive B. RNA was precipitated and bound to the spin column, contaminants were washed away with 3 serial washes, and the purified RNA was eluted in 40 µl of elution buffer. The final RNA extract was stored at -80°C until use. Small RNA libraries were prepared from 150 ng of total RNA using the Small RNA Library Prep Kit for Illumina (Qiagen), according to the manufacturer’s instructions. Briefly, 3’ adapters were ligated to the RNA and excess adapters were removed using the included column-based cleanup. Following cleanup, 5’ adapters were ligated; the input RNA was flanked by 3’ and 5’ adapters which were used to reverse transcribe the RNA. Following reverse transcription, unique indices were added through 14 rounds of PCR amplification. The final indexed PCR product was cleaned up using the included column-based cleanup. The eluate was run on a 6% TBE gel for size selection. The expected library size is ∼140bp and the corresponding band was excised from the gel and columns purified. The final libraries were quantified using a Qubit DNA High Sensitivity kit (ThermoFisher) and the size was determined using a 2100 Bioanalyzer High Sensitivity DNA Kit (Agilent Technologies, CA, USA). Normalized libraries were pooled (48 samples per pool), denatured, diluted to 0.8 pmol/l, and loaded onto a High Output (75 cycle) flow cell (Illumina, CA, USA), followed by sequencing (1 × 75 bp) on a NextSeq 550 (Illumina) ^63^.

The preprocessing of raw sequencing reads entailed removing adapter sequences and low-quality bases. Differential gene expression analysis was then conducted using edgeR within the CLC Genomics Workbench (CLCGWB) was integrated into the R/Bioconductor framework. Target enrichment and functional analysis were assessed for significant differentially expressed miRNAs, using MIENTURNET ^64^. Briefly, comparisons with at least 10 differentially expressed miRNAs were analyzed referencing the miRTarBase database of miRNA interactions. Significant interactions were defined as having an FDR adjusted p-value < 0.05 and 2 minimum interactions. The targets of the top 10 significantly enriched miRNAs were assessed for their participation in cellular processes and functions using the Database for Annotation, Visualization, and Integrated Discovery (DAVID).

### Mass Spectrometry

Proteomic mass spectrometry was performed at in the Proteomics Core Facility of the Genome Center, University of California, Davis. Briefly, peptides were resolved on a Thermo Scientific Dionex UltiMate 3000 RSLC system using An Easy Map C18 column. Separation was performed in a total run time of 90min with a flow rate of 0.520 µL/min with mobile phases A: water/0.1% formic acid and B: 80%ACN/0.1% formic acid. Peptides were analyzed on an Orbitrap Fusion Lumos (Thermo Fisher Scientific) mass spectrometer. The mass spectrometer was operated in a data-dependent acquisition mode. A survey full scan MS (from m/z 375 to 1600) was acquired in the Orbitrap at a resolution of 60k (at 200 m/z) and for MS/MS at a resolution of 15k. The LCMS files were processed with Proteome Discoverer 2.4 (Thermo Fisher) using the integrated SEQUEST engine. Precursor mass tolerance was set to 10 ppm. Fragment ion tolerance was 0.6 Da Trypsin was specified as protease, and a maximum of two missed cleavages was allowed. All data was searched against a target/decoy FASTA obtained from Uniprot. The false discovery rate of 1% was set at the PSM and protein level. To analyze the proteomic data, the STRING database (https://string-db.org/) ^65^ was utilized for protein-protein interaction networks, including both physical as well as functional associations.

### Transient middle cerebral artery occlusion (MCAO) injury

The Sprague-Dawley rats (3-4 months old) used in this experiment were purchased from Charles River Breeding Company (Wilmington, MA). They were subjected to transient MCAO by placement of a 4-0 nylon suture with a silicone coated tip into the origin of the middle cerebral artery (MCA) ^4,66^. Briefly, rats were anesthetized with nitrous oxide (70%), oxygen (30%) and isoflurane (0.5-1.5%). The right common carotid artery (CCA), external carotid artery (ECA), and internal carotid artery (ICA) were isolated. The CCA was temporarily ligated, and the branches of the ECA were dissected and divided. A 4-0 nylon suture with a silicone coated tip of 0.28 mm in diameter was advanced through the ECA and up the ICA until the tip occluded the origin of the MCA. Cerebral blood flow was monitored in the ipsilateral hemisphere by a laser-Doppler flowmeter (Moor Instruments, Devon, England, U.K.) and successful occlusion was defined by ≥ 75% decrease in baseline flow. Ninety minutes later, the suture was removed to allow reperfusion, which was confirmed by laser-Doppler flowmetry, and the ECA was ligated. Rectal temperature was kept at 37 ± 0.5^°^C during surgical procedure^1–3^. The physiological variables including mean arterial blood pressure, arterial pH, PO_2_, and PCO_2_ were measured before and after MCAO. Using this model of transient MCAO, we performed the procedure on more than 50 animals with an early mortality ∼ 10%. During the study, animals received intravenous injections of eEPC-exosomes, nh-UCBSCs, or vehicle (normal saline). Neurologic function was evaluated in all animals using a stepping test and the Neurological Severity Score (NSS) modified from De Ryck ^9^. All animals were evaluated with the NSS at 24 hours after stroke, and then in most studies, this was repeated along with the vertical pole test at 1-, 2-, 3-, and 4-weeks post stroke.

### 2,3,5-Triphenyl Tetrazolium Chloride (TTC) staining

A 1% 2,3,5-triphenyltetrzolium chloride (TTC) solution (Bd Difco) was prewarmed to 37°C in a culture plate incubator before use. Whole brains were carefully dissected and incubated at -20°C on the caps of conical tubes to facilitate tissue handling. A cutting matrix and blades were chilled with PBS prior to use. After freezing, brains were positioned ventral side up on a chilled cutting matrix. Six coronal sections each 2mm thick, were obtained using pre-chilled blades to maintain tissue integrity. Each brain slice was transferred into a well of a 12- or 6-well culture plate containing prewarmed 1% TTC solution The tissue sections were incubated at 37°C for 30 minutes, flipping the slices every 10 minutes to ensure uniform staining.

### Immunocytochemistry

For detection of NeuN, Iba1, GFAP and CD31, cells were washed once with PBS and fixed in 4% paraformaldehyde for 30–60 min at room temperature (RT). Fixed cells were washed three times with PBS, permeabilized, and blocked by incubation in blocking buffer (5% normal donkey serum, 1% BSA and 0.1% Triton X-100 in PBS) for 1 hour at RT. The cells were then incubated overnight at 4 °C with NeuN mouse (Abcam, cat# 104224), GFAP rabbit (Lifespan Biosciences, cat# B285), Iba1 goat (Abcam, cat# 5076, 1:500), MAP2 (Invitrogen, cat# PA1-16751), VEGFR2 (Abcam, cat# ab2349), CD31 (Abcam, cat# ab64543) antibodies with Alexa Fluor 488 at a dilution of 1:500 in 5% normal donkey serum, 0.5% BSA and 0.05% Triton X-100 in PBS. Then, the cells were washed four times with PBS plus 0.05% Triton X-100 (PBS-Triton) and incubated for 1 hour at RT with secondary Abs at a 1:1000 dilution in PBS-Triton. Isotype controls were stained under identical conditions, except that total rabbit IgG (Life Technologies, 10500C) was used as the primary antibody. Cells were counterstained with DAPI at 0.1 ng/mL for 10 min at RT and imaged on a Nikon Eclipse TE2000-U inverted microscope.

### Data processing and statistical analyses

All data are expressed as mean ± standard deviation. Statistical analysis between groups at each time point were performed by the unpaired student’s t-test. Independent experiments of samples over time were analyzed by repeated measures of ANOVA with the Holm adjustment. Differences were considered significant at probability values of P < 0.05.

## Results

### Purification of exosomes from human embryonic endothelial progenitor cells (eEPCs)

We previously established human eEPC lines that display functional properties of endothelial cells as well as embryonic gene expression patterns. The eEPCs were shown to be highly scalable, having up to 80 population doublings (pd) and stable long-term expansion of over 50 pd with stable angiogenic properties at late passage ^58^. Importantly, eEPC-derived exosomes enhanced tube formation *in vitro,* and their angiogenic activity was retained during scale-up in a Quantum bioreactor ^58^. In this study, we developed standardized and quantitative exosome purification methods that are rapid, cost-effective, reproducible and scalable with an aim toward pre-clinical and clinical applications.

For exosome purification, we collected supernatant after incubating eEPCs for 72 hours at 5% oxygen in exosome-free conditioned medium. To develop the exosome purification protocol, we assessed two methods of tangential flow filtration (TFF) and size exclusion chromatography (SEC) (**Figure 1A-B**) and compared an SEC-only isolation method to a combination method of TFF and SEC. TFF is a simple and scalable filtration method based on molecular weight cut off (MWCO) size exclusion to separate exosomes from impurities (e.g., proteins) ^67^. Size exclusion chromatography (SEC) is a gravity alone or low-speed technology using a column containing porous beads which separates according to molecular size ^68^. First, to verify the biological activity of exosomes isolated from the two methods, we tested angiogenic activities of exosomes derived from three independent eEPC lines (**Suppl. Figure 1A-B**). We found that SEC exosome separation preserved the biological function of exosomes but resulted in a low yield (**Figure 1C**). The combination of tangential flow filtration (TFF) and size exclusion chromatography (SEC) achieved a ∼ 100-fold increase in purity (1E10-5E10 particles/ug of protein) and a yield of eEPC-exosomes that was ∼10-fold higher than SEC alone (**Figure 1C**). This purity and yield of exosome purification meets ISEV consortium guidelines ^46,47^ for identifying highly purified exosomes, which allowed us to scale up production. Importantly, the combined TFF and SEC method retained or increased angiogenic potency of eEPC-exosomes as indicated by endothelial wound-healing assay (**Suppl. Figure 1A**) and live-cell tube formation assay (**Suppl. Figure 1B**). This result showed that the combined methods of TFF and SEC could achieve production of highly purified exosomes for mass production by mainlining biological potency.

**Figure 1.**
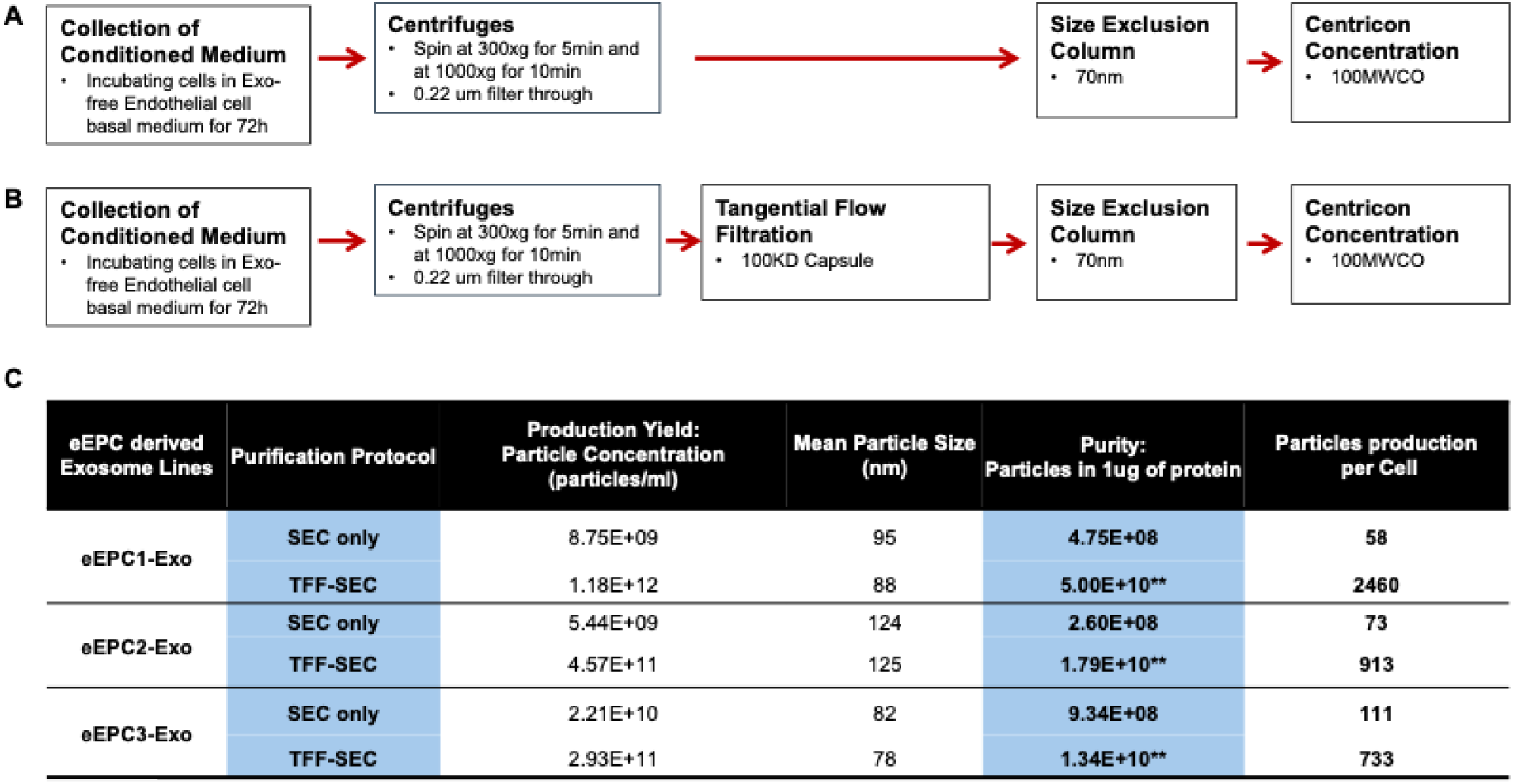
The development of the exosome purification protocol. Methods isolating eEPC-exosomes using (A) size exclusion chromatography (SEC) only and (B) combining tangential flow filtration (TFF) and a size exclusion chromatography (SEC). (C) Analysis results by comparing SEC only versus TFF-SEC combined scaling-up exosome purification methods (** exosome purification meets 2024 ISEV consortium guidelines).

### Characterization of human eEPC-exosomes

To assess the characteristics of exosomes isolated using the established protocol combining TFF and SEC methods, three eEPC-derived exosome lines (eEPC1-exo, eEPC2-exo and eEPC3-exo) were tested. Using immunoblot analysis, we identified the expression of exosome surface markers such as CD63, CD81, ICAM, EpCAM, and TSG101 (**Figure 2A**), but no cellular contamination marker, GM130 (Cis-Golgi matrix protein), was detected. Exosome particles were analyzed by both tunable resistive pulse sensing (TRPS) and nanoparticle tracking analyses (NTA), which validated size distribution, concentration, and purity (**Figures 2B-C**). The mean particle size of eEPC-exo lines were approximately 80-125nm (eEPC1-exo, 88nm; eEPC2-exo, 125nm and eEPC3-exo, 78nm). Protein concentration was determined using a micro-BCA assay, and a particle-to-protein ratio of no less that 1E10 particles per 1µg of protein was set as a minimum accepted purity (**Figure 1C**) ^69^. These results indicated that eEPC-exosomes that were purified using combined methods of TFF and SEC were free of detectable cellular contamination.

**Figure 2.**
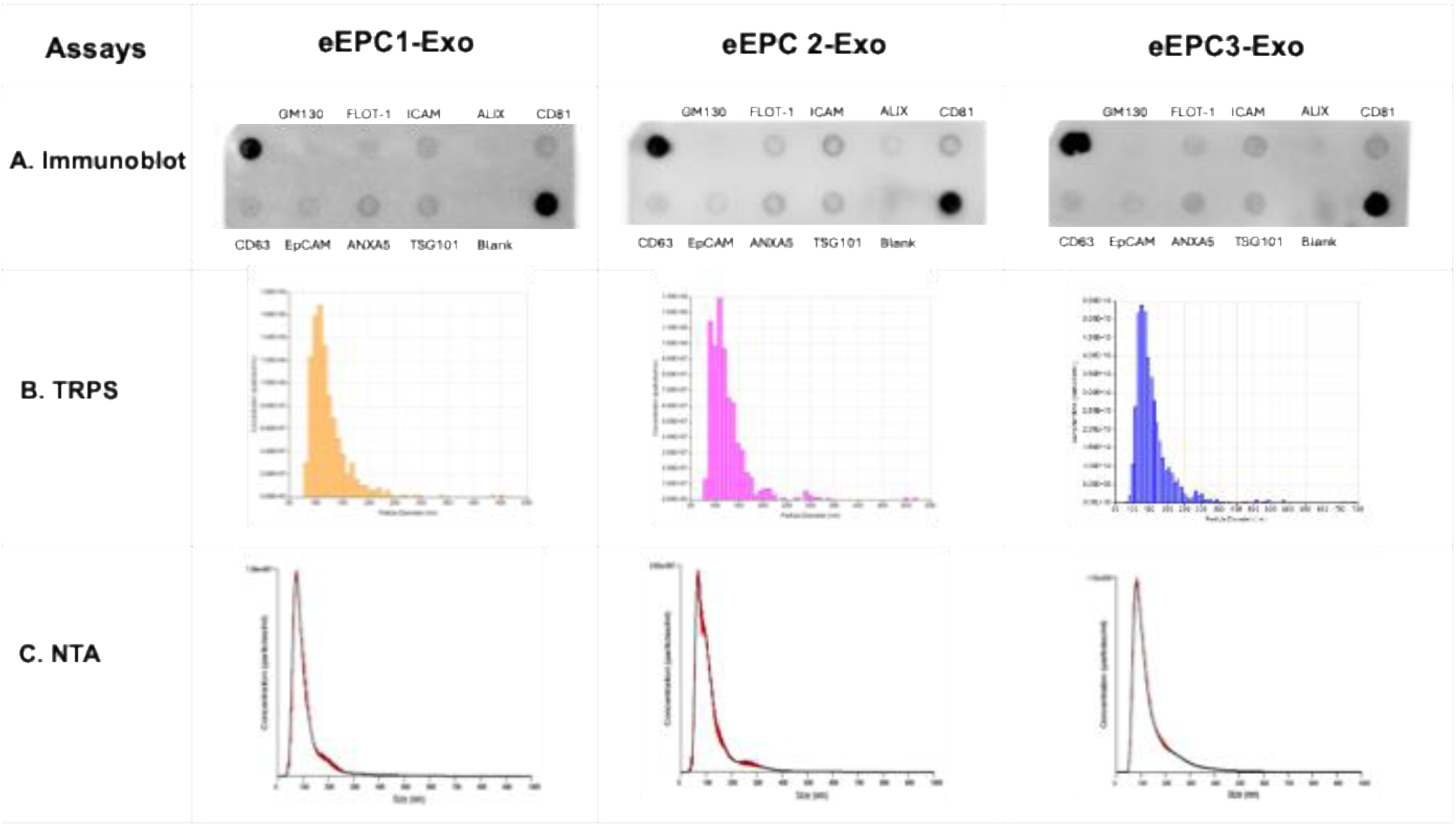
Characterization of three eEPC-derived exosome lines (eEPC1-exo, eEPC2-exo and eEPC3-exo). (A) immunoblot assay to identify the protein expression of exosome surface markers and particles analysis using (B) Tunable Resistive Pulse Sensing (TRPS) as well as confirmed with (C) nanoparticle tracking analysis (NTA). The mean particle size of eEPC-exo lines were approximately 80-120nm (eEPC1-exo, 88nm; eEPC2-exo, 125nm and eEPC3-exo, 78nm).

### Assessing the angiogenic activity of human eEPC-exosomes

We next investigated *in vitro* bioactivity of eEPC-exo lines using live-cell imaging assays that measured endothelial wound-healing and tube formation activity (**Figure 2**). To assess wound healing migration activity of eEPC-exosomes, three doses of particles (1E2, 1E3, and 1E4 particles per cell) were tested to human umbilical vein cells (HUVECs), comparing to VEGF treatment **(Suppl Figure 2A)**. Exosome depleted medium (Exo-Free-MV2) was used for baseline control. HUVECs were monitored every 2 hours for 24 hours. We found that eEPC-exo lines were directly taken up by HUVECs and promoted cell migration at concentrations as low as 1E3 exosomes/cell with no increase or inhibition of activity seen at higher doses **(Suppl Figure 2A)**. Next, we compared the angiogenic activity of eEPC-exosomes with growth factor VEGF (10ng/ml) and MSC-derived exosomes. Interestingly, both eEPC-exosomes resulted in faster scratch wound healing activity starting from 6 hours to 16 hours of migration (p>0.001) than primary MSC-derived exosomes and an equivalent migration activity to VEGF (10ng/ml) (**Figure 3A** and **Supplementary Video 1**). These data suggest that eEPC-derived exosomes contain higher angiogenic potency, which involves ‘accelerated’ migration and/or replication of the target HUVEC cells compared to MSC-derived exosomes.

**Figure 3.**
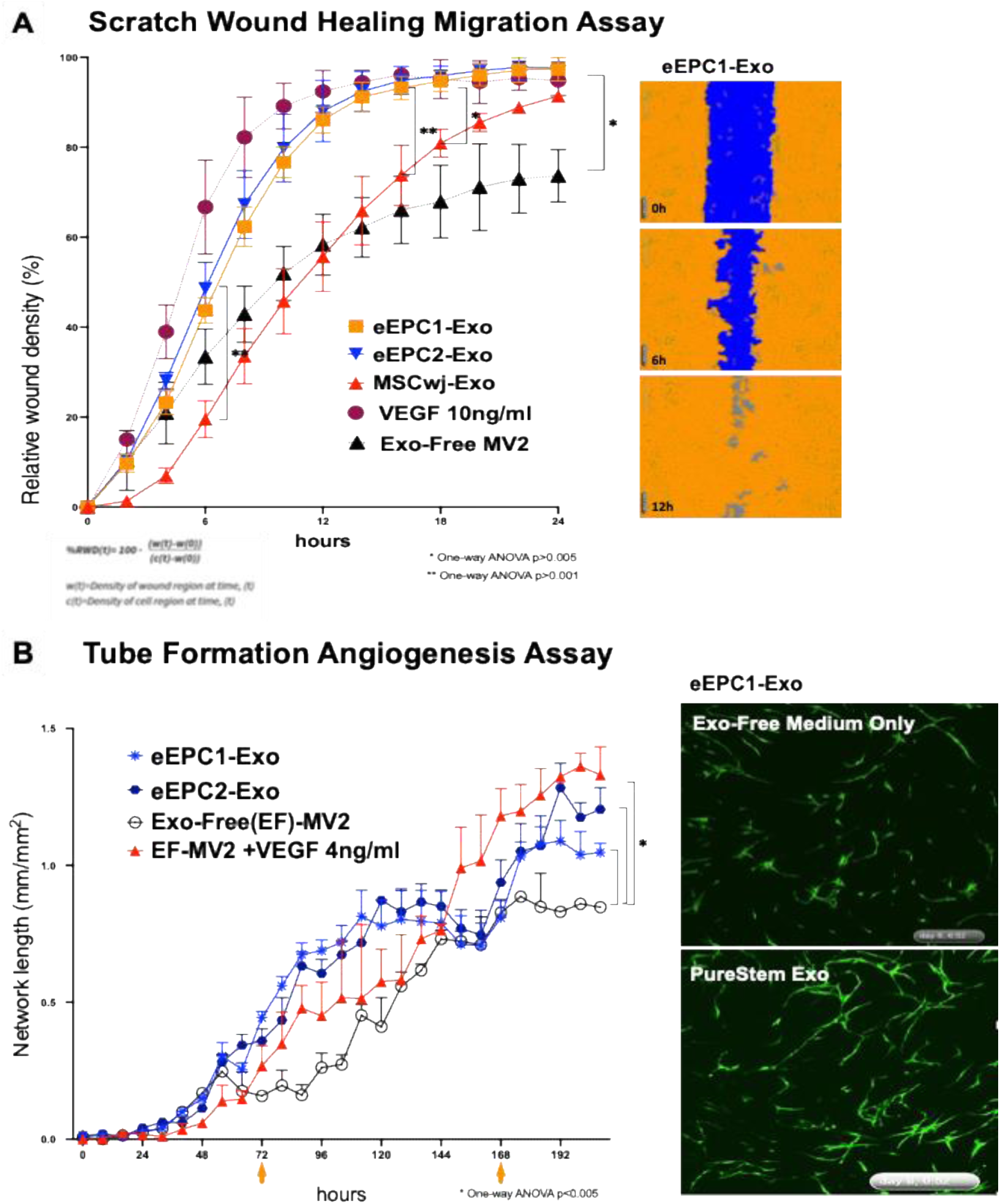
Validating in vitro angiogenic bioactivity of eEPC-Exosomes. **(A)** Scratch wound healing assay in HUVECs monitored every 2 hours for 24 hours using IncuCyte ZOOM. Migration data were analyzed as the Percent of Relative Wound Density (% RWD). **(B)** Tube formation angiogenesis assay using IncuCyte 96-Well Kinetic Angiogenesis PrimeKit. Progenitor exosomes were directly taken up by HUVECs and promoted the migration, proliferation, and tube formation of endothelial cells. We found that progenitor exosomes showed higher angiogenic potency than primary MSC-derived exosomes. Average Network Length (mm/mm^2^): Average of the length of all the networks in the image divided by the image area (mm^2^). (One-way ANOVA *p > 0.005, **p > 0.001).

Additionally, we verified the distinct bioactivity of eEPC-exosomes comparing to fibroblasts derived exosomes (**Suppl Figure 2B)**, showing that eEPC-exosomes resulted in faster scratch wound healing activity than fibroblast-exosomes. It is indicated that the effect of exosomes can be as potent as that of parent cells ^58^ in promoting angiogenesis. When we induced the inactivation of possible exosome carriers such as RNAs and proteins, using heat and protease K treatment, the promoted migration activity by the treatment of eEPC-exo was abolished without damaging the cells as level of basal medium cultured cells (**Suppl Figure 2C)**. It is implicated that the bioactivity of exosomes was due to the biological properties of exosome cargo.

Furthermore, we confirmed the angiogenic functionality of eEPC-exosomes using live cell quantitative analysis of HUVEC endothelial cell tube formation angiogenesis assay (**Figure 3B** and **Supplementary Video 2**). To assess tube network growth of HUVECs treated with eEPC-exosomes, HUVECs were labeled using TagGFP and co-cultured with normal human dermal fibroblasts (NHDF). This model allowed us to demonstrate all phases of the angiogenesis process, including proliferation, migration, and, eventually, differentiation and formation of vascular networks. Imaging the co-culture in a live-cell analysis system enabled us to distinguish GFP-labeled HUVECs from co-cultured NHDF and to visualize the vessel formation networks of HUVECs over 8 days **(Figure 3B and Supplementary Video S2**). eEPC-exosomes were directly treated to cells at day 3 and day 7 and tube formation in HUVECs was monitored every 4 hours over 8 days. To quantify the amount of tube formation, we combined time-lapse image acquisition to measure network tube length. Tube formation over the 8 days was quantified as average network length using an angiogenesis analysis module according to the manufacturer (Sartorious AG). **Figure 2B** shows eEPC1-exo and eEPC2-exo stimulated HUVEC tube forming activity that was comparable to VEGF (4 ng/ml), where an integral proangiogenic cytokine is shown as the positive control. These data demonstrate the angiogenic activity of eEPC-exo, further demonstrating their angiogenic and proliferative potency.

### Determination of miRNA cargo composition of human eEPC-exosomes

To further characterize eEPC-exosomes, we first validated miRNA cargo using RNA-seq analysis. Small RNA-sequencing analysis revealed that three eEPC-exosome lines (eEPC1-exo, eEPC2-exo, and eEPC3-exo) enriched miRNAs related to mediating angiogenesis, controlling inflammation, and remodeling tissue (**Supple Table 1**). Notably, eEPC2-exos (which had the highest wound healing activity) lack miR-126 but contain miR-210 and miR29-b/c, which are absent in the other EPC-exos analyzed. The eEPC-exo lines were significantly enriched in miR126, miR-92, miR130, miR-221, and miR132 relating to angiogenesis, miR-192 and miR-29 relating to remodeling. miR155, miR-21, miR146 relating to controlling inflammation, and miR-21-5p relating to anti-apoptosis (**Supple Table 1**). In particular, the eEPC1-exosomes contained six angiogenic miRNAs including miR-126-3p, miR-192, miR-92a, miR130, miR-221/222, and miR132 (**Figure 4A).** Using MIENTURNET (MicroRNA ENrichment TURned NETwork), we input a list of top 50 miRNAs and analyzed miRNA-target interactions. First, the reactome analysis of the six angiogenic miRNAs revealed that these miRNAs regulate interleukin signaling, VEGF pathway, and MAPK signaling cascade (**Figure 3B)**. Next, Gene Ontology (GO) pathway enrichment analysis was performed using 249 target genes of the top 50 most abundant miRNAs in eEPC1-exosomes and was identified as enriching gene targets involved in vascular development, neurogenesis, and brain development (**Figure 3C).** We also investigated GO biological pathway analysis of 50 abundant miRNAs’ target genes in eEPC2-exosomes and found that nervous system development, immune system development, vascular/blood development, and cell adhesion pathways are enriched (**Supple Figure 3)**. Our data show that eEPC1-exosomes contain high levels of angiogenic miRNAs cargos (i.e., miR-126, miR-192, miR-92, miR130, miR-221, miR132), anti-inflammatory miRNA (i.e., miR-146, miR-155, miR-21-5p and miR-146), and anti-apoptotic miRNAs (i.e., miR-21-5p), suggesting that eEPC1-exo could deliver miRNA cargo and modulate target genes, which improves functional recovery, enhances neurogenesis, and inhibits neuroinflammation in an ischemic stroke model.

**Figure 4.**
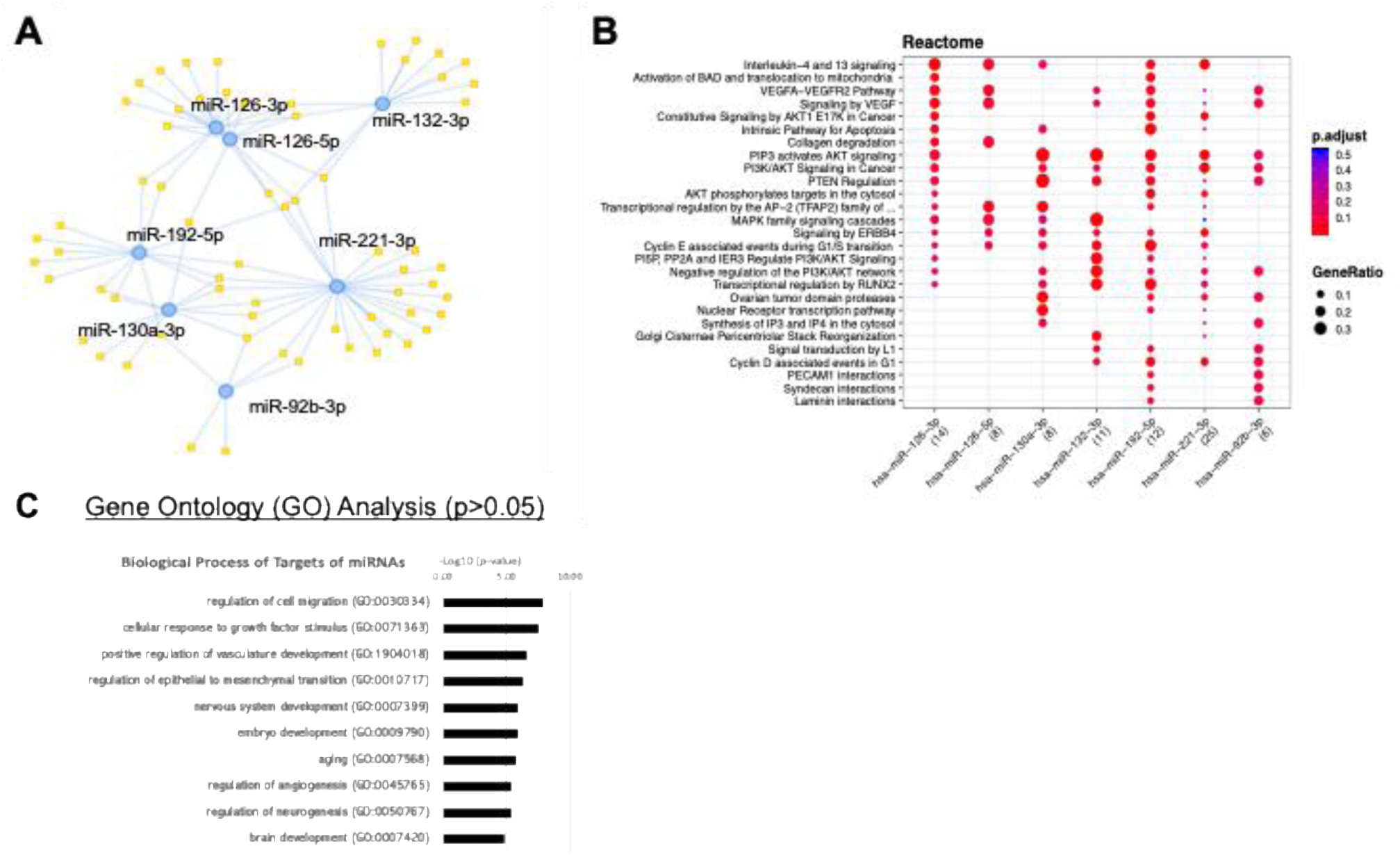
Small RNA sequencing analysis in eEPC1-Exosomes. (A) eEPC1-exosomes express a set of angiogenic miRNAs (miR-126, miR-192, miR-92, miR130, miR-221, miR132), (B) Reactome pathway analysis of 6 angiogenic miRNAs, **(C)** GO pathway enrichment analysis of top 50 miRNA target genes using DAVID, based on the differentially expressed genes (DEGs) with p-values (false discovery rate; FDR) < 0.05 and an absolute fold change (FC) > 2.0.

### Determination of protein cargo composition of human eEPC-exosomes

Next, to determine the protein cargo of eEPC-exosomes, we performed mass spectrometry proteomic analyses. The STRING database (https://string-db.org/) ^65^ was utilized to enhance the functional enrichment analysis of cargo proteins in eEPC-exosomes. This allowed us systematically to collect and integrate protein-protein interaction networks, including both physical as well as functional associations. We analyzed the top 27 most abundant proteins in eEPC1-exosomes (**Figure 5 and Suppl Table 2**) and the top 36 abundant proteins in eEPC2-exosomes (**Suppl Figure 4 and Suppl Table 2**). The abundant 27 protein cargos in eEPC1-exosomes were clustered with 4 groups based on physical association to each other in a protein complex or in a transient complex (**Figure 5A)**. In parallel with the network prediction, we assigned the 27 proteins to their corresponding pathways using Gene Ontology (GO) biological process and found that eEPC1-exosomes contained a group of proteins displaying functions that include epidermis Development (DSP; FABP5; CDSN; CASP14), astrocyte development (VIM; S100A9), and aging (IGFBP2; EEF2; CAT), as well as promotion of cell migration (KRT6A; DSP) (**Figure 4B)**. The top 36 proteins in eEPC2-exosomes were clustered in 5 groups, and the proteome included proteins associated with biological cell adhesion, cell migration, cell migration, and wound healing, including VAMP3, FAP, ITGAV, ICAM1, and ANXA5 (**Suppl Figure 4A-C and Suppl Table 2**).

**Figure 5.**
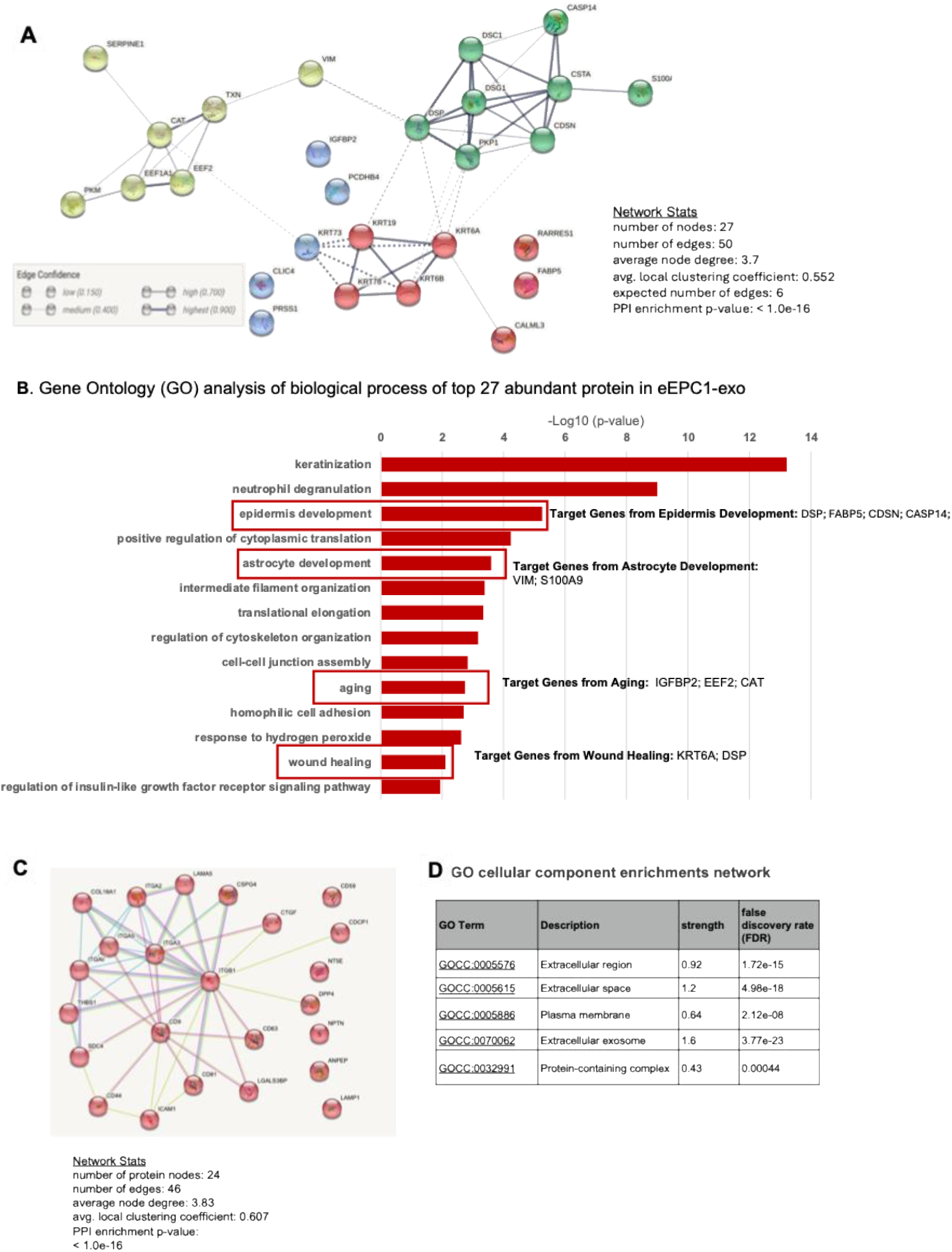
Mass spectrometry protein analysis in eEPC1-exosomes. (A) Interaction network of top 27 abundant proteins in eEPC1-exosomes, (B) Gene ontology analysis of biological process, (C) extracellular vesicle component protein interaction network in eEPC1-exosomes, (D) GO Cellular component enrichments network of 46 extracellular vesicle component proteins

Additionally, we validated that the total proteomes of eEPC-exosomes are highly enriched for known extracellular vesicular proteins. Consistent with our previous data, we detected signatures of a typical set of proteins associated with the exosome including CD9, CD81, and CD63 (**Figure 5C-D**). GO cellular component enrichment analysis data showed that these proteins were related to extracellular plasma proteins, exosomes, and protein-containing complex.

In the process of selecting angiogenic exosome candidates for application to the stroke animal model, we ensured the quality control and reproducibility of exosome production. Regarding cargo contents including miRNAs and proteins, we found differential characteristics between eEPC1-Exo and eEPC2-Exo, but similarities between eEPC1-Exo and eEPC3-Exo. We recently reported the characterization of clonal progenitor cell lines including eEPC1, eEPC2, and eEPC3, indicating that these eEPC lines contain angiogenic capacity, but show distinct gene expression as endothelial progenitors (EPC1 and EPC3) and perivascular progenitors (EPC2) ^58,70^. Due to the different cell type origin of each exosome production cell line, we found a difference among the purified exosomes from these lines in terms of particle size (**Figure 2**), exosome protein expression (Figure 2), and cargo contents (**Suppl Table 1** and **Suppl Table 2**). Cargo analysis data also indicate that there are two distinct types of exosome lines between eEPC1-exo and eEPC2-exo, but also that they contain common miRNA molecular targets as well as biological enrichment pathways, such as wound healing. We expect this result will provide more options to investigate the effective therapeutic exosomes of stroke therapy in future animal model applications. In the current report, we chose the eEPC1-exo production line as a source of exosomes to test a preclinical stroke model because of its significant angiogenic potential and miRNA content, including miRNA-126 cargo ^71^. In future studies, it would be interesting to test eEPC2-exosomes both alone and in combination with eEPC1-exosomes in the treatment of stroke.

### Intravenous delivery of eEPC1-exosomes promotes the recovery of neurological function in rat MCAO model

We tested the therapeutic feasibility of eEPC1-exosomes using a rat MCAO stroke model. eEPC1-exosomes at 3E10 particles were administrated intravenously to ischemic MCAO rats after 24 hours of stroke onset. A modified neurological severity score (mNSS) that shows sensorimotor function was compared at 1, 7, 14, 21, and 28 days after stroke **(Figure 6A)**. We found that administration of eEPC1-exosomes (Blue line) significantly improved limb placement motor function 7 days following MCAO as compared to the saline group (Red line), which showed no improvement at day 7 and resulted in near complete recovery in PureStem-exosome treated animals **(Figure 6D-E)**. Interestingly, eEPC1-exosomes showed higher neurological improvement at day 7 (\**p* < 0.05) and day 14 (\*\**p* < 0.01), which involves ‘*accelerated’* recovery compared to UCBSC treated animals (Green line) based on mNSS score **(Figure 6D)**. Next, we observed that eEPC1-exosomes highly improved performance on the vertical pole **(Figure 6E)**. The velocity of each animal’s descent down a gridded vertical pole was calculated and compared across groups over time. After a stroke, rats tend to perform poorly on this behavioral task due to asynchronization of gripping ability on the affected side. In contrast, administration of eEPC1-exosomes significantly improved the change in velocity at 7, 14, and 21 days after stroke compared to the saline group (**Figure 6D-E**). Furthermore, we observed reduced inflammatory response at the histological level in eEPC1-exosome treated animals after MCAO. **Figure 7** shows that rats with eEPC1-exosome treatment had a significant reduction in the number of GFAP-positive and IBA1-positive cells (glia and astrocyte markers, respectively, \**p* < 0.05) in the right motor cortex (stroke infarct) region as compared to the saline control group. The neuronal marker, NeuN, increased in eEPC1-exosome treated animals after MCAO (\**p* < 0.01), providing evidence that exosomes may reduce inflammatory cascade events in ischemic brain injury and increase neuronal survival. Taken together, the results showed that eEPC1-exosomes highly improved the recovery of neurological function in a rat MCAO model, suggesting eEPC-exosomes play a role in inducing neurogenesis and modulating inflammation in ischemic injury.

**Figure 6.**
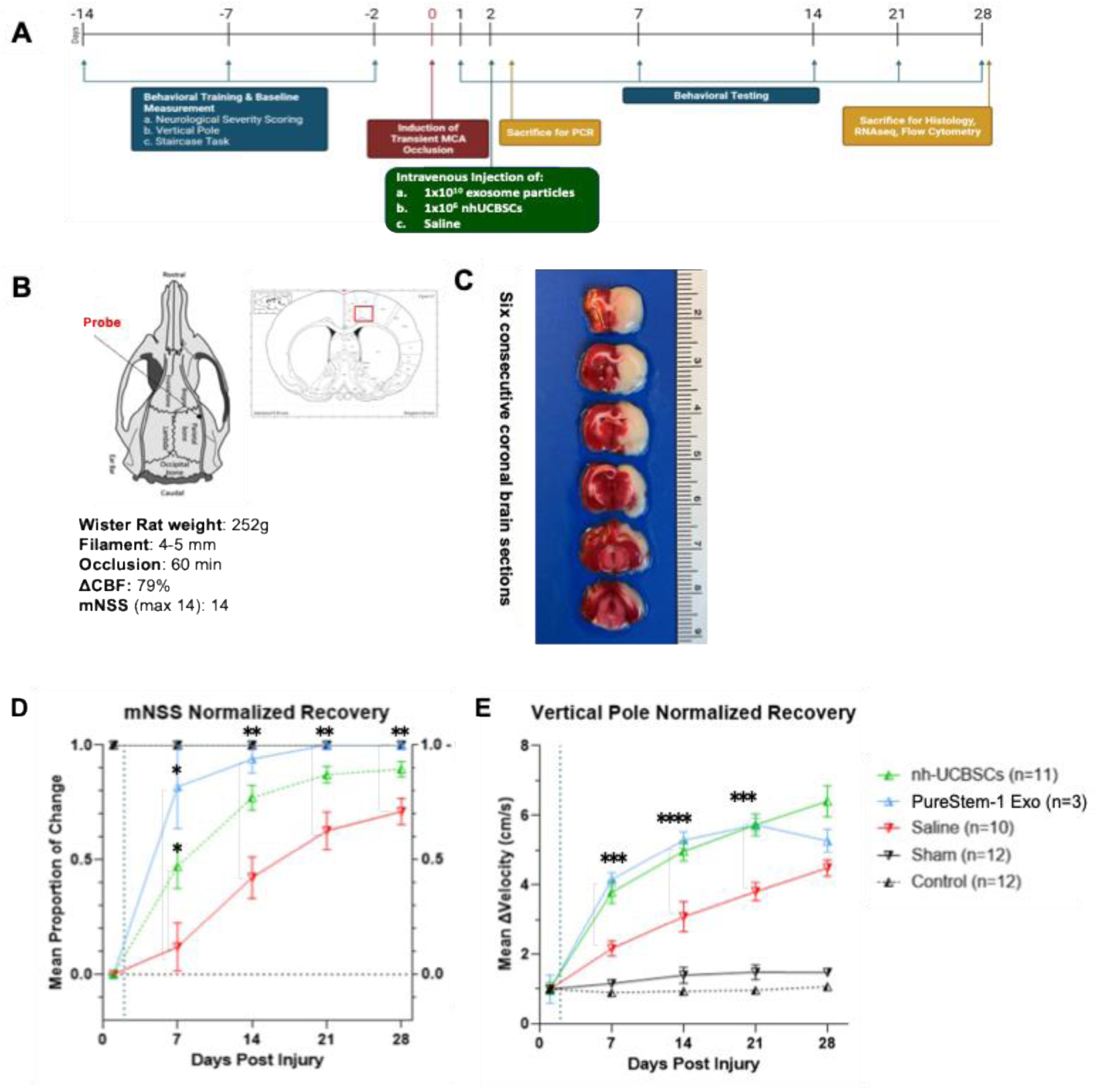
Demonstration of Preclinical Therapeutic Potential in rat Middle Cerebral Artery Occlusion (MCAO) Animal Model. (A) Experimental design and study criteria of MCAO model, (B) Scheme of an intraluminal suture MCAO model, (C) Six consecutive coronal brain sections after transient MCAO. (D) Modified neurological severity score (mNSS) and (E) vertical pole shows normalized recovery test, which measures sensorimotor function was significantly increased by intravenous administration of eEPC1 exosomes at 3E10 particles in ischemic MCAO rats. A Two-way Analysis of Variance (ANOVA) with Tukey’s Multiple Comparison Test was performed for statistical analysis (*p < 0.05 and **p < 0.01).

**Figure 7.**
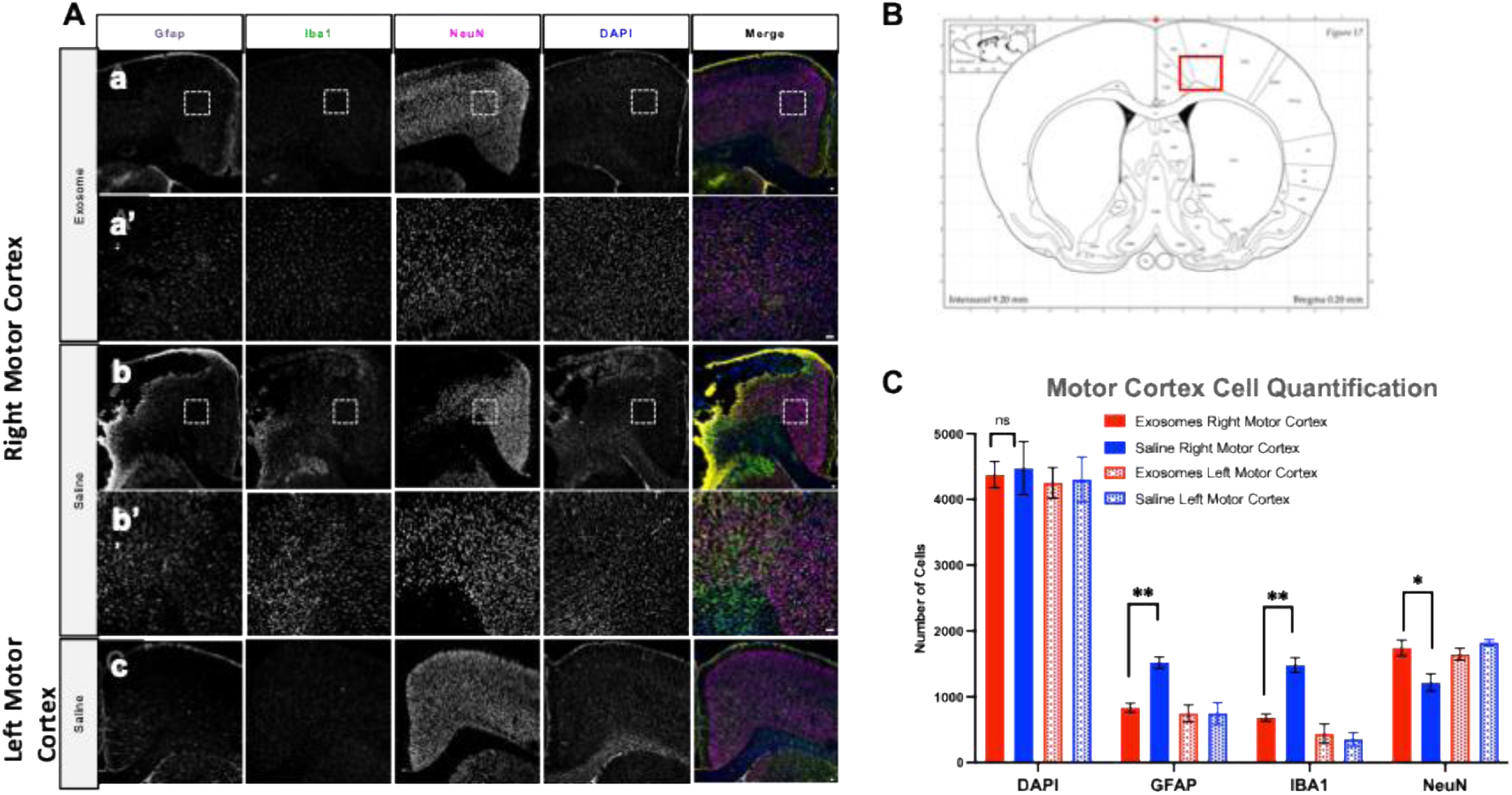
Immunofluorescent staining of (A) GFAP, astrocyte marker; Iba1, microglial marker; NeuN, neuronal marker; and DAPI, nuclear DNA. (a-c) 4x magnification; (a’ and b’) 10x magnification and Scalebar = 100um, (B) Representative coronal maps showing stroke infarct legion in right motor cortex. (C) Motor cortex cell quantification in eEPC1-exosome treated and saline treated rats, (One-way ANOVA *p > 0.005, **p > 0.001).

## Discussion

In this report, we have detailed the development of an optimized exosome purification methodology using human embryonic endothelial progenitor cell (eEPC) lines as a stable and scalable source of exosome production. In particular, we found that eEPC-derived exosomes (eEPC-exosomes) contain angiogenic, neurogenic, and immunomodulatory cargo and are active in scratch wound and vascular network tube forming assays indicating angiogenic activity. eEPC-exosomes contain high levels of miRNA cargos related to angiogenesis, anti-inflammation, and anti-apoptosis. Furthermore, eEPC-exosomes contain a group of proteins related to astrocyte development and aging, as well as promotion of cell migration. These data support the idea of that the systemic administration of eEPC-exosomes might improve functional recovery after ischemic stroke by stimulating angiogenesis, preventing inflammatory cascade events and increasing neuronal regeneration. Thus, we tested the therapeutic efficacy of one eEPC-exosome candidate (eEPC1-exo) in rat model of ischemic stroke model (MCAO). Our *in vivo* data demonstrate the potential efficacy of eEPC-exosomes for reducing neurological deficits when administered systemically at 24 hours after onset of stroke.

Exosomes are defined by their small size (30-120nm), the presence of certain transpanins (CD63, CD81, CD9), and secretion via multivesicular bodies. They are secreted by most cell types and play a key role in inter-cellular communication through the transfer of their cargo of lipids, proteins, and RNAs to recipient cells ^72^. The therapeutic effects of exosomes are mainly attributed to their powerful ability to transfer molecular cargos (i.e., miRNAs and proteins), which reduces secondary injury and stimulates natural tissue repair mechanisms. Multiple studies have evaluated the effects of exosomes derived from primary stem cells in ischemic disease including acute ischemic stroke (MSCs, NPCs, and BMSCs) ^19,30,73,74^ and myocardial infarction (EPCs and MSCs) ^75,76^, demonstrating promising therapeutic effects through the promotion of angiogenesis, alleviation of inflammatory response, and improvement of functional neurologic recovery. However, a significant bottleneck impeding translation of primary stem cell-derived exosomes to clinical development lies in the limitations of manufacturing platforms for exosome production. Inherent limitations of primary stem cells include low proliferative capacity, donor variability, and population heterogeneity, all of which hinder industrial scale exosome production for clinical application toward treating disease ^5,6^. Moreover, our current and previous data ^58^ indicate that embryonic progenitor exosomes have increased potency over adult stem cell exosomes (i.e., MSC). This is likely because of their origin in the highly regenerative embryonic stage of development. In this report, we investigated the novel application of eEPC-exosomes using a highly scalable and clonally pure eEPC lines platform ^58^ as a source of exosomes for stroke therapy ^57,58^.

For stroke therapy, current evidence suggests that exosome treatments will be well tolerated and have a wide therapeutic window of days rather than hours ^35,38,77^. As such, exosome therapies represent a potential paradigm shift in the management of stroke by overcoming the existing dogma that stroke intervention beyond 12 hours is of no benefit. Importantly, for systemic stroke treatment, exosomes can cross the blood brain barrier (BBB) ^78,79^. Previously, we demonstrated the efficacy of using human UCBSC-derived stem cells in treating rats with MCAO ^16,80^ and neonatal hemorrhagic brain injury ^81^. In the present study, we have demonstrated that intravenous administration of eEPC-exosomes to ischemic rats substantially enhances angiogenesis, anti-inflammatory neuroprotection, and behavior improvement. Our data show that exosomes not only carry functional cargos, which facilitate angiogenesis, but also protect against neuroinflammation in stroke. eEPC1-exosomes contain high levels of angiogenic miRNAs cargos (i.e., miR-126, miR-192, miR-92, miR130, miR-221, miR132), anti-inflammatory miRNA (i.e., miR-146, miR-155, miR-21-5p and miR-146), and anti-apoptotic miRNAs (i.e., miR-21-5p). miR-126 is one of the most abundant microRNAs in eEPC1-exosomes and plays a crucial role in regulating the function of angiogenesis and vascular integrity ^82,83^, as well as embryonic angiogenesis ^60,72^. Emerging studies have demonstrated that intravenous administration of overexpressed miR-126 exosomes post-stroke improved functional recovery, proving that miR-126 plays a key role in initiating neurorestorative mechanisms in the brain such as vascular remodeling, anti-inflammation, and neurogenesis ^82,84, 83^. We also identified angiogenic and scratch wound healing activity of eEPC-exo lines (i.e., eEPC1-exo, eEPC2-exo and eEPC3-exo). eEPC3-exos are made from eEPC3, which is closely related to the eEPC1 production line ^58^. eEPC2-exos appear to be a variation of eEPC1-exos but contain lower levels of miR-126. The eEPC2-exos and their production line are distinct from eEPC1 and eEPC3 ^58^. We detected clear differences in miRNA cargo, including the absence of miR-126 and presence of two additional miRNAs (i.e., miR-210 and miR-29b/c) not present in the other two exosomes (**Supple Table 1**). Thus, the angiogenic and scratch wound activity may be the result of a distinct mechanism of action. Of note is the increased activity of eEPC2-exos in the scratch wound assay compared to the other exosomes. Further studies are planned to explore the therapeutic value of eEPC2-exos alone or in combination therapy with EPC1-exos for treating stroke and other degenerative conditions.

Our proteomic mass spectrometry analysis study found that eEPC1-exosomes carry a group of proteins related to astrocyte development (VIM; S100A9) and aging (IGFBP2; EEF2; CAT), as well as promote cell migration (KRT6A; DSP). These data are consistent with the higher angiogenic activity of eEPC1-exosomes and suggest their potential to improve functional outcomes in the treatment of ischemic stroke. Indeed, we observed significant improvement in neurological and cognitive function seen at 14 to 21 days post-ischemic injury under treatment with eEPC1-exosomes, which may be attributed to downstream signaling pathways and neurorestorative mechanisms. The precise mechanisms underlying facilitation of angiogenesis and neurogenesis due to administrated exosomes will require further studies. However, it is unlikely that a single exosome mediator such as growth factors or microRNAs, which also induce neurogenesis or angiogenesis, is responsible for these combined effects ^20,61,83,85^. Rather, we hypothesize that there is a complex interplay between multiple cargo components of exosomes, which has yet to be fully elucidated. Many recent studies have documented that transplanted stem cells and their secreted exosomes, such as neural progenitor cells and mesenchymal stem cells, are able to modify the post-stroke microenvironment in terms of growth factor contents and enhanced post-stroke neurogenesis ^21,30,35,83,86^. Based on these observations, we conclude that eEPC-exosomes can alleviate ischemic injury and promote revascularization and neurological function recovery in an MCAO model, demonstrating the preclinical efficacy of eEPC-exosomes in stroke therapy.

In terms of exosome purification methodology, so far, the most commonly used exosomes isolation methods in the field (i.e. ultracentrifugation, chemical precipitation, ultrafiltration etc.) have resulted in impurities in the precipitate, including protein aggregates, virion, sub-cellular organelles, and damaged exosomes, which may cause a reduction in their biological activity ^87^. Here, we have developed a scalable and stable method that can provide intact and pure exosomes. As such, we believe our approach is pivotal in developing an ideal exosome purification methodology, in particular, one where producing subsequent downstream therapeutic molecules such as exosome cargo components are effective. Toward this end, we developed methods that combine TFF and SEC as a simple and economical purification method for handing large-scale GMP compliant exosome production for preclinical and clinical exosome therapy.

Recent preclinical and clinical investigations suggest that exosomes released by stem cells may recapitulate the therapeutic effects of their donor cells without the drawbacks inherent in stem cell therapy ^53^. Indeed, recent reports have shown that exosomes account for much of the therapeutic effects with no apparent undesired off-target effects. Dendritic cell-derived exosomes and ascites-derived exosomes have been administered directly and are shown to be safe, non-toxic, and well tolerated when used for cancer immunotherapy in clinical trials ^88,89^. Moreover, multiple doses of MSC-exosomes have been administered to a patient with GvHD with no side effects ^90^. In contrast with stem cells, exosomes are non-replicative, so they can eliminate the risk of unwanted cell growth. Another significant advantage of exosomes is their low immunogenicity. Our results indicate a lack of MHC class I and II antigens on eEPC-produced exosomes (data not shown), which could help them avoid immune surveillance ^91^. Thus, we believe a key advantage of using eEPC-exosomes for stroke therapy lies in their potential as an allogeneic treatment with robust stability, which enables development of an off-the-shelf product for stroke intervention. Together, these features make exosomes much more practical, safe, and convenient than cell therapy. Developing a standardized protocol for manufacturing therapeutic exosomes that expand uniformly on a large scale with control over quality will be critical in paving the way for human clinical studies using exosome therapy for stroke.

## Conclusion

The present study has established an exosome purification methodology that achieves robust and scalable exosome production using novel clonally pure embryonic endothelial progenitor cell lines and demonstrated the potential of eEPC-exosomes as a therapeutic for alleviating ischemic stroke injury and inflammation. This work will enable further preclinical developments toward the use of novel exosome stoke therapies, which can ultimately support studies for treating human stroke.

## Research ethics statement

Animal experiments were conducted under the protocol approved by the Institutional Animal Care and Use Committee (IACUC), Animal Welfare Assurance Number A3456-01 and the National Institutes of Health Guide for the Care and Use of Laboratory Animals (NIH Publications No. 80-23, revised 1996.). To avoid or minimize pain and distress, animals were monitored daily by a combination of Animal Care technicians, veterinary technicians, and individual laboratory staff. Veterinary consults were requested for animals that show unusual or unexpected behaviors.

## Author Contributions

Conceptualization, D.L., J.L. and A.G.; methodology, J.L., D.C., M.H., and S.V.; software, J.L.; validation, J.L. and D.L.; formal analysis, J.L., D. N-R., D.C., S.V.; investigation, J.L. and A.G., W.C.L., D.L.; resources, A.G., and M.D.W.; writing—original draft preparation, J.L. and D.L.; writing—review and editing, J.L., D.L., A.G., and W.C.L.; visualization, J.L.; supervision, J.L. and A.G; project administration, J.L., D.L. and A.G.; funding acquisition, J.L., D.L. and A.G. All authors have read and agreed to the published version of the manuscript.

## Data Availability Statement

Data is available upon request.

## Acknowledgements

We thank Dr. Gabriela Grigorean for performing LC-MS/MS and data analysis in the Proteomics Core Facility of the Genome Center, University of California, Davis and Drs. Paul Robbins and Fernando Santiago at University of Minnesota for guiding RNAseq data analysis.

## Patents

Exosomes from Clonal Progenitor Cells, 2022 (US 11274281); Exosomes from Clonal Progenitor Cells, 2019 (US 10,240,127); Methods and compositions for targeting progenitor cell lines, 2019 (US 10,227,561).

---

**Suppl Fig 1.**
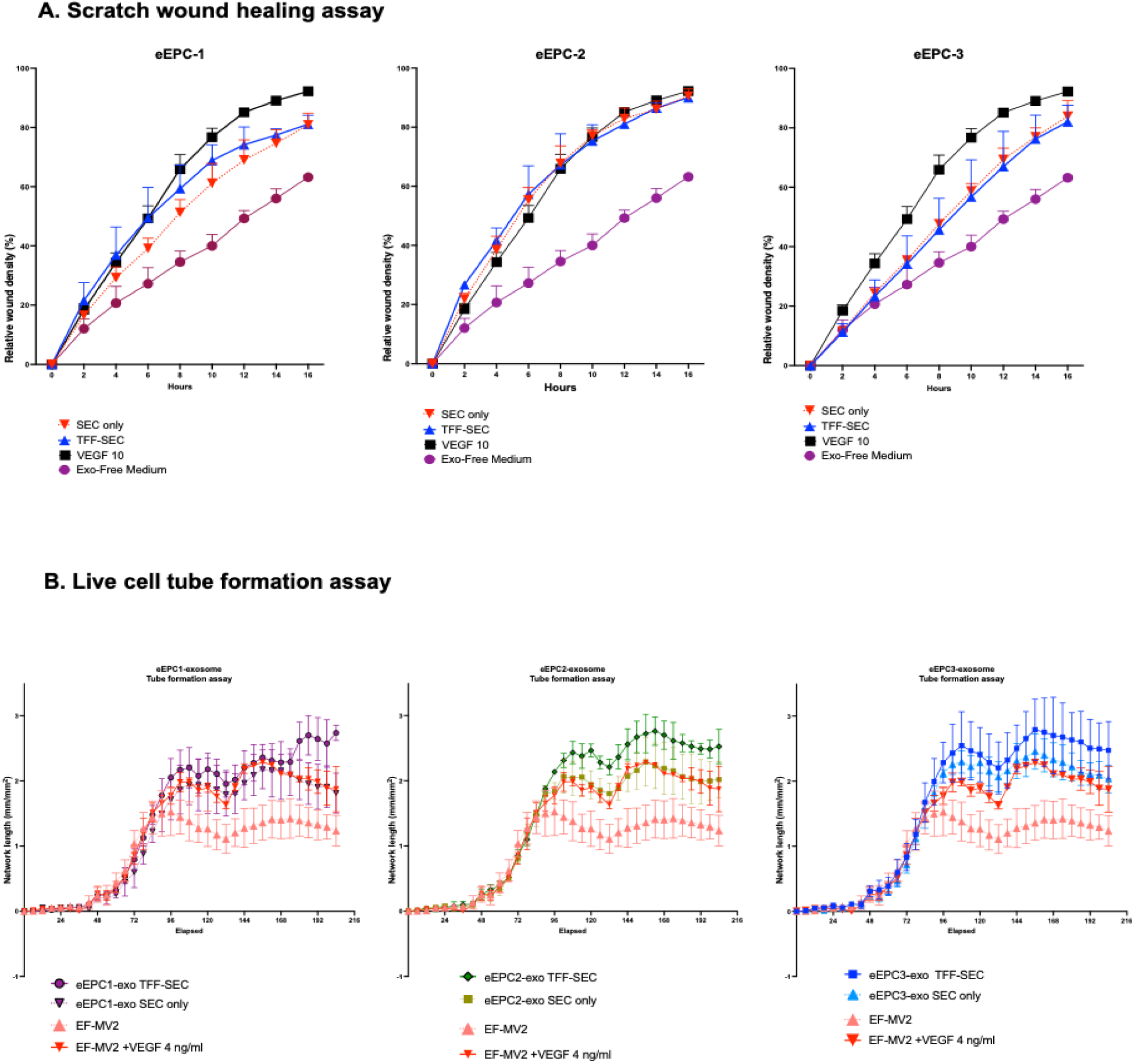
Comparison of functional activity in three eEPC exosome lines (eEPC1-exo, eEPC2-exo and eEPC3-exo) isolated from TFF-SEC vs. SEC only, (A) Scratch wound healing assay and (B) Live cell tube formation assay

**Suppl Fig 2.**
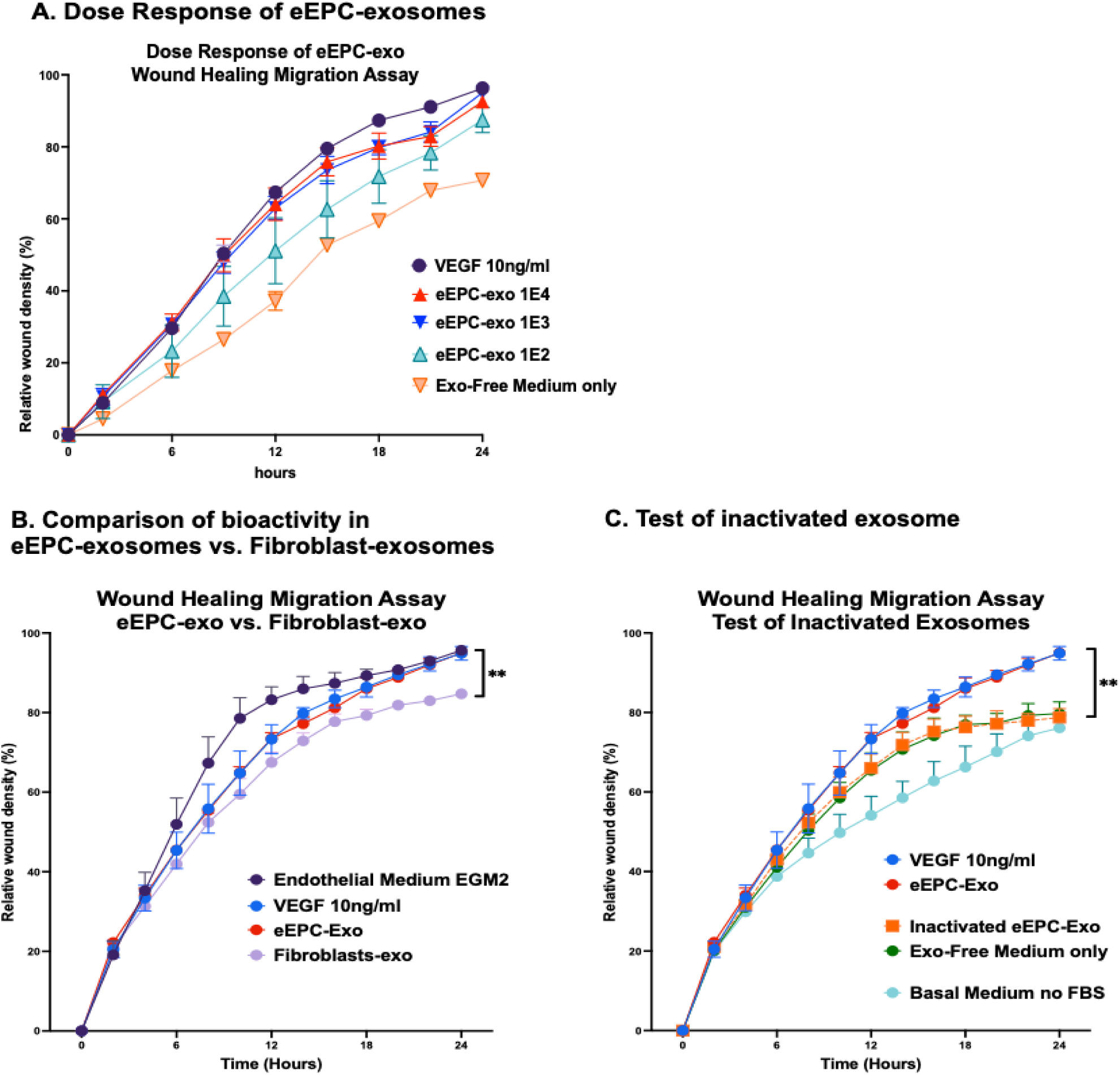
Migration assays for (A) Dose Response of eEPC-exosomes, (B) Comparison of bioactivity in eEPC-exosomes vs. Fibroblast-exosomes, and (C) Test of inactivated exosome.

**Supple Fig 3.**
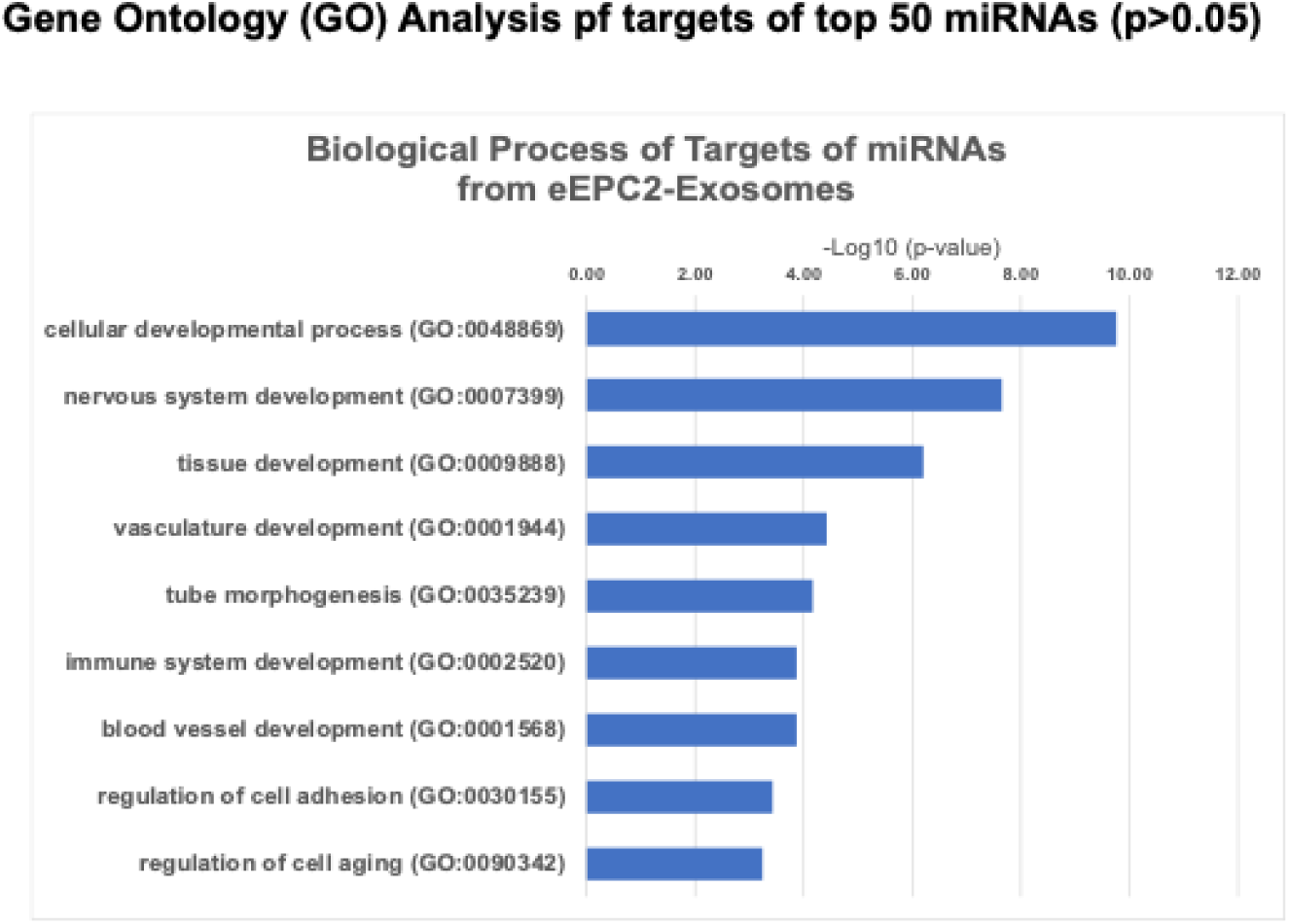
Gene Ontology Analysis of eEPC2-Exosome Small RNA-seq.

**Suppl Figure 4.**
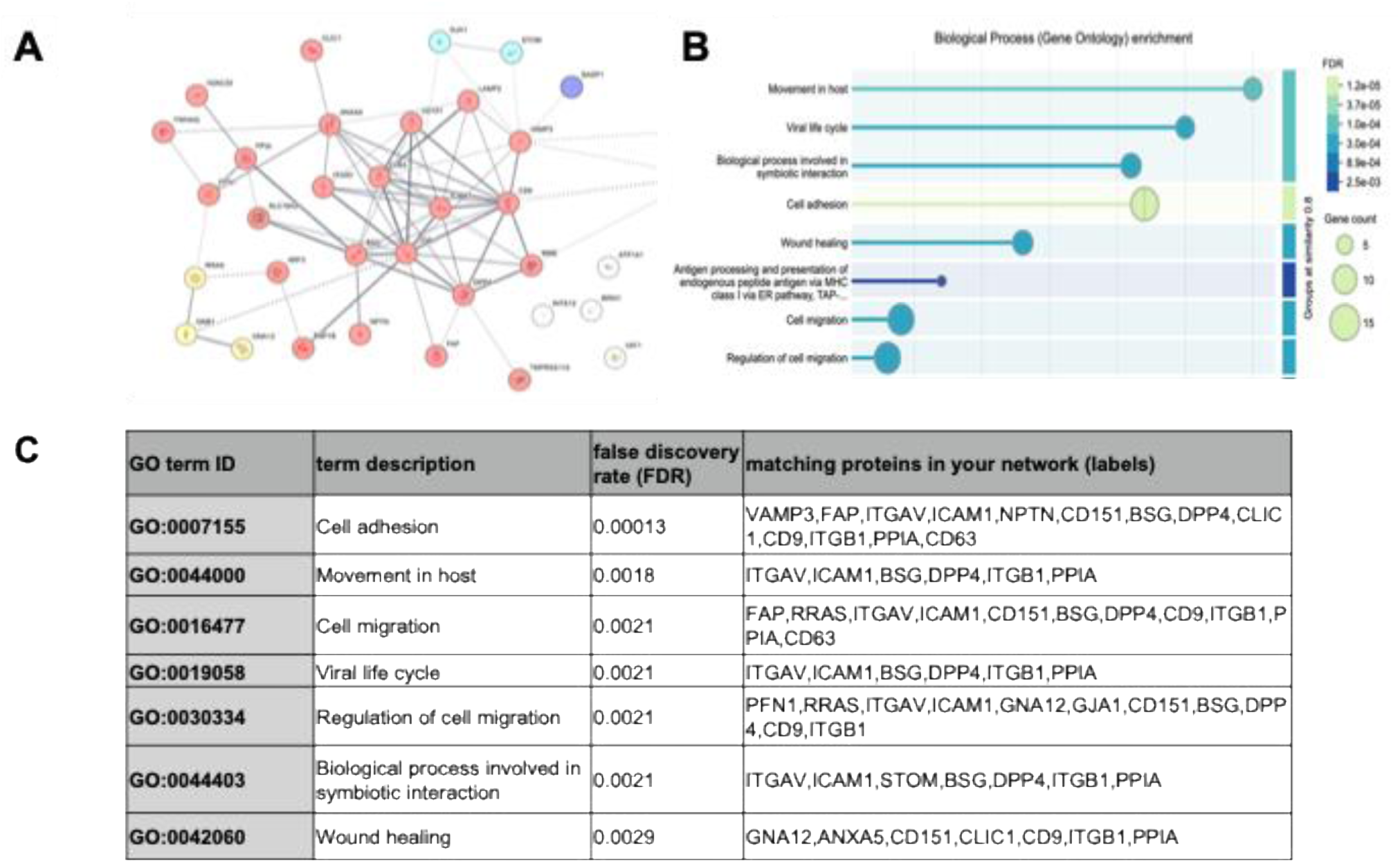
Mass spectrometry protein analysis of top 36 abundant protein in eEPC2-exosomes. (A) Interaction network of top 36 proteins in eEPC2-exosomes, (B) Gene ontology analysis of biological process in eEPC2-exo, and (C) List of proteins enrichment network.

**Suppl Table 1.** List of most abundant miRNAs from small RNAseq analysis in eEPC-exo lines.

|  | eEPC1-exo | eEPC2-exo | eEPC3-exo | miRNA target(s) | Function |
| --- | --- | --- | --- | --- | --- |
| hsa-miR-126 | ✓ | x | ✓ | Spred-1 | EC-mediated angiogenesis |
| hsa-miR-192 | ✓ | ✓ | ✓ | SIP1 | Remodeling |
| hsa-miR-92a | ✓ | ✓ | ✓ | ITGB5 | EC-mediated angiogenesis |
| hsa-miR-130 | ✓ | ✓ | ✓ | GAX, HOXA5 | EC-mediated angiogenesis |
| hsa-miR-132 | ✓ | ✓ | ✓ | p120RasGAP | EC-mediated angiogenesis |
| hsa-miR-221/222 | ✓ | ✓ | ✓ | c-kit, eNOS*, p27/Kip1 | EC-mediated angiogenesis |
| hsa-miR-155 | ✓ | ✓ | ✓ | SHIP1, SOCS1, IL12 | Wound Healing: Control of inflammation |
| hsa-miR-20a | ✓ | ✓ | ✓ | VEGF, E2F1 | Wound Healing: EC-mediated angiogenesis |
| hsa-miR-17-5p | ✓ | ✓ | ✓ | TIMP1 | EC-mediated angiogenesis |
| hsa-miR-20a | ✓ | ✓ | ✓ | E2F1 | EC-mediated and tumor induced angiogenesis |
| hsa-miR-378 | ✓ | ✓ | ✓ | Sufu, Fus-1 | Angiogenesis |
| hsa-miR-155 | ✓ | ✓ | ✓ | AT1R | EC-mediated angiogenesis |
| hsa-miR-210 | x | ✓ | x | ISCU1/2, E2F3 | Re-epithelization, EC-mediated and tumor induced angiogenesis |
| hsa-miR-29b/c | x | ✓ | x | Smads, beta-catenin | Wound Healing: Remodeling |
| hsa-miR-21-5p | ✓ | ✓ | ✓ | PTEN, PDCD4 | Control of inflammation and apoptosis |
| hsa-miR-146 | ✓ | ✓ | ✓ | IPAK, COX2 | Control of inflammation |

**Suppl Table 2.**
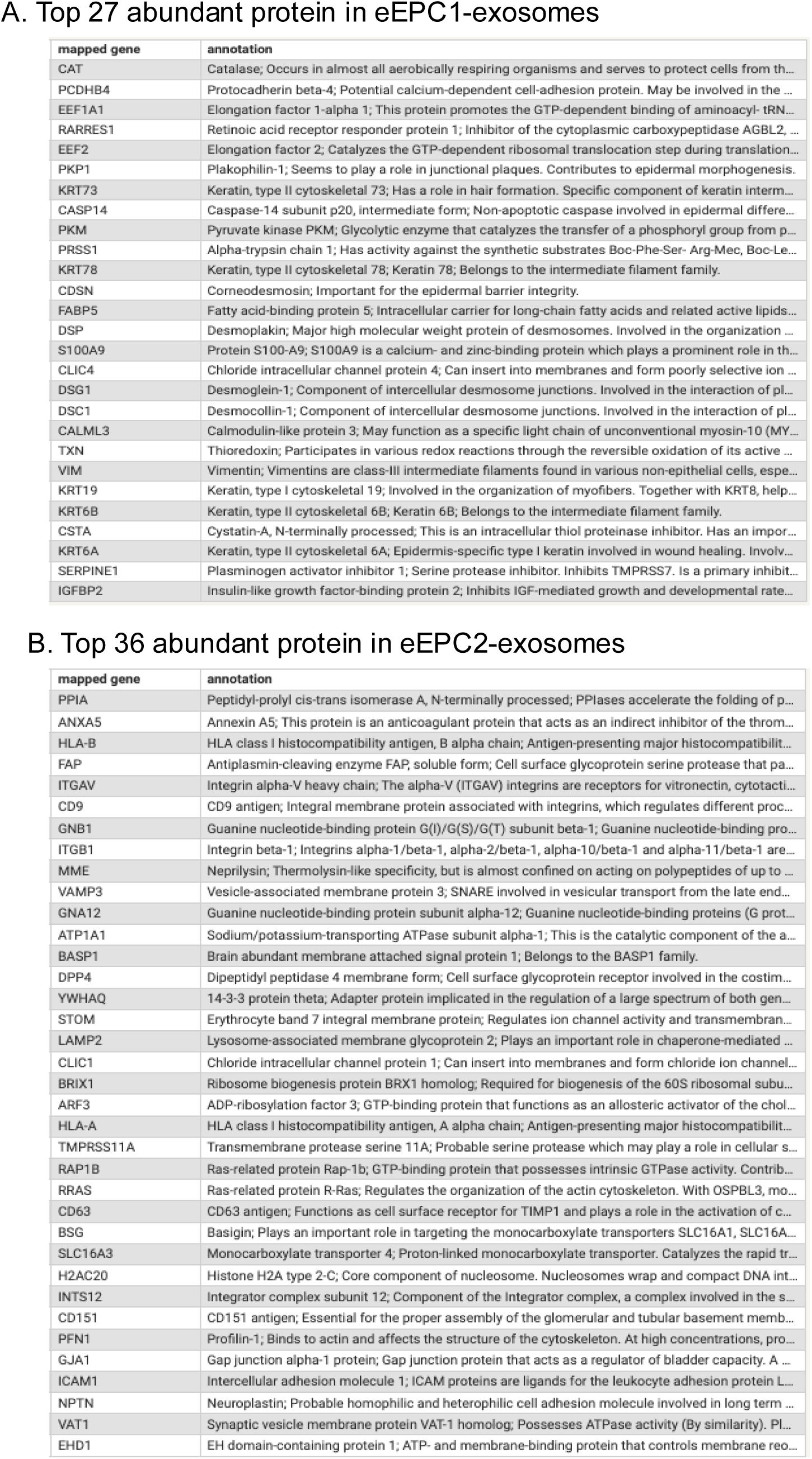
Mass spectrometry protein analysis list of (A) top 27 abundant protein in eEPC1-exosomes and (B) top 36 abundant protein in eEPC2-exosomes.

